# Spatial organization of single mRNPs at different stages of the gene expression pathway

**DOI:** 10.1101/237008

**Authors:** Srivathsan Adivarahan, Nathan Livingston, Beth Nicholson, Samir Rahman, Bin Wu, Olivia Rissland, Daniel Zenklusen

## Abstract

mRNAs form ribonucleoprotein complexes (mRNPs) by association with proteins that are crucial for mRNA metabolism. While the mRNP proteome has been well characterized, little is known about mRNP organization. Using a single molecule approach, we show that mRNA conformation changes depending on its cellular localization and translational state. Compared to nuclear mRNPs and lncRNPs, association with ribosomes decompacts individual mRNAs, while their sequestration into stress-granules leads to increased compaction. Moreover, translating mRNAs rarely show co-localizing 5’ and 3’ ends, indicating that mRNAs are either not translated in a closed-loop configuration, or that mRNA circularization is transient, suggesting that a stable closed-loop conformation is not a universal state for all translating mRNAs.

**One Sentence Summary:** Single mRNA studies in cells show RNA compaction changes depending on translational state, but mRNAs are not translated in closed-loop conformation.

---

mRNAs are single-stranded nucleic acid polymers. Intramolecular base pairing and binding of RNA binding proteins (RBPs), many of which contain homo and hetero-dimerization domains, assemble mRNAs into mRNPs (*1*). Yet how mRNPs are organized as three-dimensional particles within cells remains unknown (*2–10*). mRNPs may exist as compact assemblies or as flexible open polymers allowing frequent interactions between different regions required to regulate mRNA metabolism at different stages of the gene expression pathway, such as during translation or RNA turnover (*11–13*). To study mRNP organization within cells, we combined Structured Illumination Microscopy (SIM) with single molecule resolution fluorescent in situ hybridization (smFISH) and Gaussian fitting to investigate the spatial relationship of various regions within mRNAs in different cellular compartments and translational states.

To determine if this approach allows to spatially resolve different regions within single mRNAs, we first measured co-localization precision by hybridizing alternating probes, labeled with cy3 and cy5, to a 1.2 kb region within the 18,413 nt-long MDN1 mRNA in paraformaldehyde-fixed HEK293 cells. Images were acquired spanning the entire cell volume, and 3D datasets reduced to 2D by maximum intensity projection. Fig. 1A shows co-localization of signals emitted from both channels detecting single MDN1 mRNAs. We determined the center of each signal by 2D Gaussian fitting and measured the distance between signals from both channels (*14, 15*). We obtained a co-localization precision of 21 nm, indicating that we can resolve discrete regions within mRNAs separated by more than 20 nm (Fig. 1C). We then positioned labeled probes to the 5’ and 3’ ends of MDN1 to determine maximal extension of the MDN1 mRNA in cells (Fig S1 and Table S1), which, fully extended, would measure 5.5 m in length. Upon analyzing cytoplasmic mRNAs, we observed few overlapping 5’ and 3’ signals; however, the majority of 5’ signals had a 3’ signal within close proximity (Fig. 1B), with distances up to 300 nm between the two signals. A similar distribution was observed when measured in 3D, and distances were indistinguishable when the EtOH step in the hybridization protocol was omitted (Fig. S2B, C). To determine if 5’ and 3’ signals were part of the same mRNA molecule, we used a third set of FISH probes tiling the entire length of the mRNA between the 5’ and 3’ regions in 500 nt intervals. The tiling signal overlapped with either one of the two regions and connected the 5’ and 3’ within the 300 nm radius, confirming that 5’ and 3’ end signals belonged to the same molecule and, moreover, pointing towards an elongated conformation of cytoplasmic MDN1 mRNPs (Fig. 1D). To better understand the spatial relationship between different regions within these mRNAs, we replaced the tiling probes with a probe set hybridizing to the middle region of the MDN1 mRNA (Fig. 1E, S1). Using these probes, we observed cytoplasmic mRNAs in various configurations in which the three different regions could be spatially resolved (Fig. 1E and F). To measure the average volume of these cytoplasmic mRNAs, we aligned individual mRNAs using their center of mass and found a mean radius of gyration (<R_g_>) of 73.95 nm (Fig. 1G).

**Figure 1:**
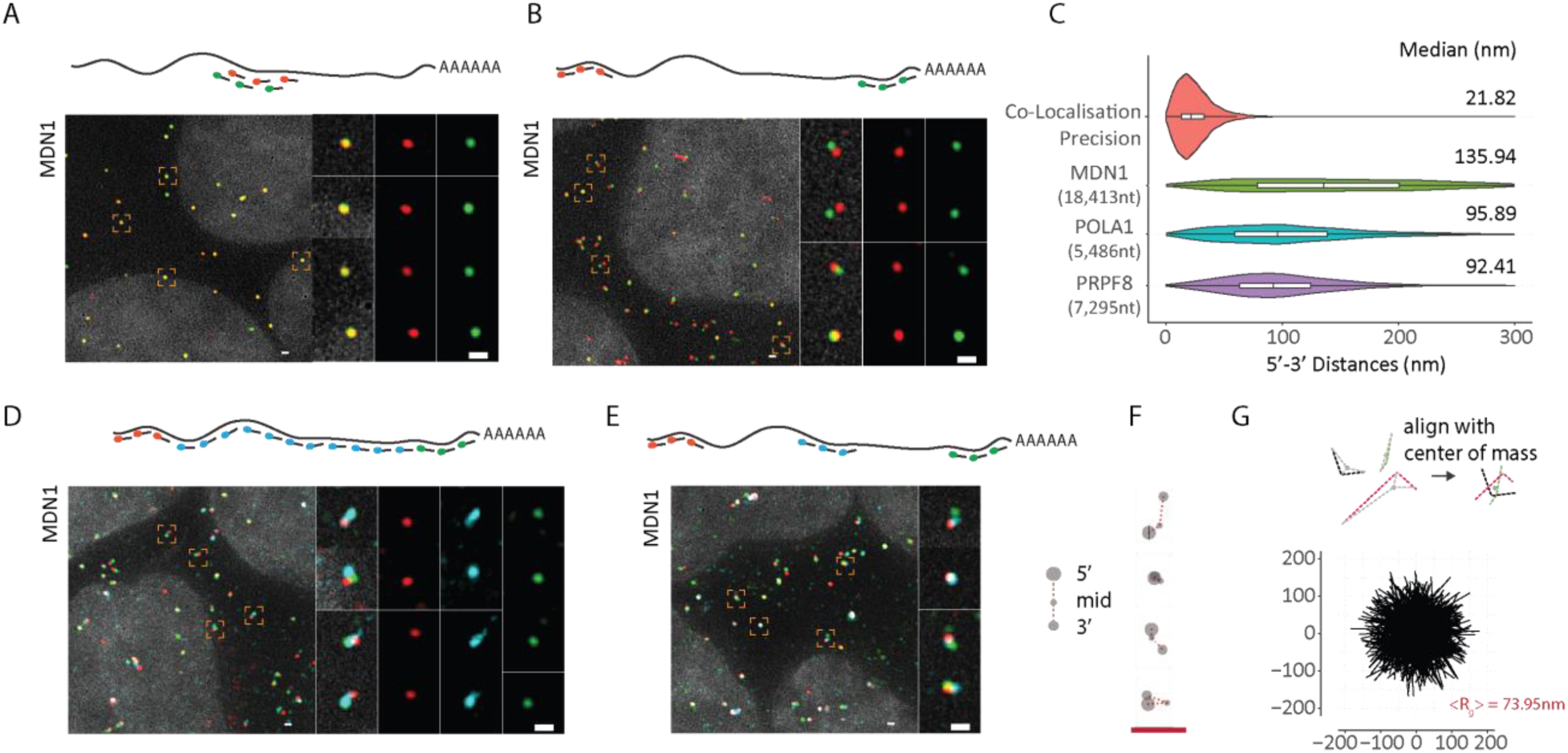
Visualizing single mRNA reveals open conformations of cytoplasmic mRNAs. **(A)** smFISH images using alternating probes labeled in cy3 (red) and cy5 (green) to middle region of MDN1 mRNA in paraformaldehyde fixed HEK 293 cell. Nuclei are visualized by DAPI staining (grey). Magnified images of individual RNAs marked by dashed squares are shown on the right. Schematic position of probes shown on top. **(B)** smFISH using probes targeting the the 5’ (red) and 3’ (green) ends of MDN1. **(C)** Violin plots showing distance distribution of co-localization precision of co-localizing spots from A, and 5’-3’ distances for MDN1, POLA1, PRPF8 mRNAs determined by Gaussian fitting. White box plot inside the violin plot shows first quartile, median and third quartile. Median distances are shown on the right. **(D, E)** smFISH using 5’ (red), 3’ (green), and tiling or middle probes (cyan) respectively. (**F)** Cartoon depicting different mRNA conformations from E. **(G)** Projections of superimposed conformations from E with their centers of mass in registry, n=563. Mean Radius of gyration (<R_g_>). Scale bars, 500 nm.

To determine whether such open conformations are particular to the long MDN1 mRNA or are a more common feature of cytoplasmic mRNAs, we measured compaction of two shorter mRNAs encoding for the splicing factor PRPF8 (7,295 nt) and the DNA polymerase alpha catalytic subunit POLA1 (5,486 nt) and found similar open conformations (Fig S3A). End-to-end distances showed narrower distributions compared to MDN1, suggesting that maximum expansion in cells is limited by mRNA length. Together, these data show that cytoplasmic mRNAs exist in an open conformation where 5’ and 3’ are rarely found in close proximity.

Translating mRNPs are thought to exist in a closed-loop conformation where 5’ and 3’ ends of the mRNA are brought together through interactions between the cap binding eIF4F complex and the polyA binding protein PABPC1 (*11, 12, 16*). Surprisingly, 5’-3’ conformations consistent with such a closed-loop configuration were rarely observed. One possibility could be that most mRNAs with separated 5’ and 3’ ends are not in the process of being translated and that only the fraction with co-localizing ends represents the pool of translating mRNAs. If that were to be the case, interfering with translation should further reduce the fraction of mRNAs with co-localizing 5’ and 3’ ends. To test this hypothesis, we treated cells, prior to fixation, with drugs that affect translation via different mechanisms: cycloheximide interferes with translation by binding to the E-site of the 60S ribosomal unit and stabilizes polysomes, whereas puromycin causes premature chain termination and disassembles polysomes (*17*). Treatment with cycloheximide only minimally affected the distribution of 5’-3’ end distances when compared to untreated cells (Fig. 2C). However, the disassembly of polysomes following a short treatment with puromycin (10 min) resulted in an unexpected phenotype where the 5’-3’ ends of most transcripts were found co-localizing (Fig. 2A). Distance measurements showed a narrow distribution with a median of 36 nm. Treatment with the translation inhibitor homoharringtonine that stalls ribosomes at the initiation site for 1 hour, yielded similar results (Fig. S2B). Moreover, POLA1 and PRPF8 ends showed a high degree of co-localization with similar median 5’-3’ end distances (Fig. 2C and S3B). This observation could either represent a change in mRNP conformation resulting in increased levels of 5’-3’ interaction, or it could be the result of a general compaction of the mRNP due to the loss of bound ribosomes. Probes hybridizing to the middle region of MDN1, or tiling probes along the entire length of the MDN1 transcript, showed that puromycin treatment resulted in a general compaction of the mRNPs (Fig. 2B, S3C). Overlaying mRNA conformations revealed a much less extended form of these mRNPs compared to untreated cells (Fig. 2D and E). These observations suggest that most cytoplasmic mRNAs are translated, that mRNAs within translating mRNPs are not arranged in a closed-loop conformation.

**Figure 2:**
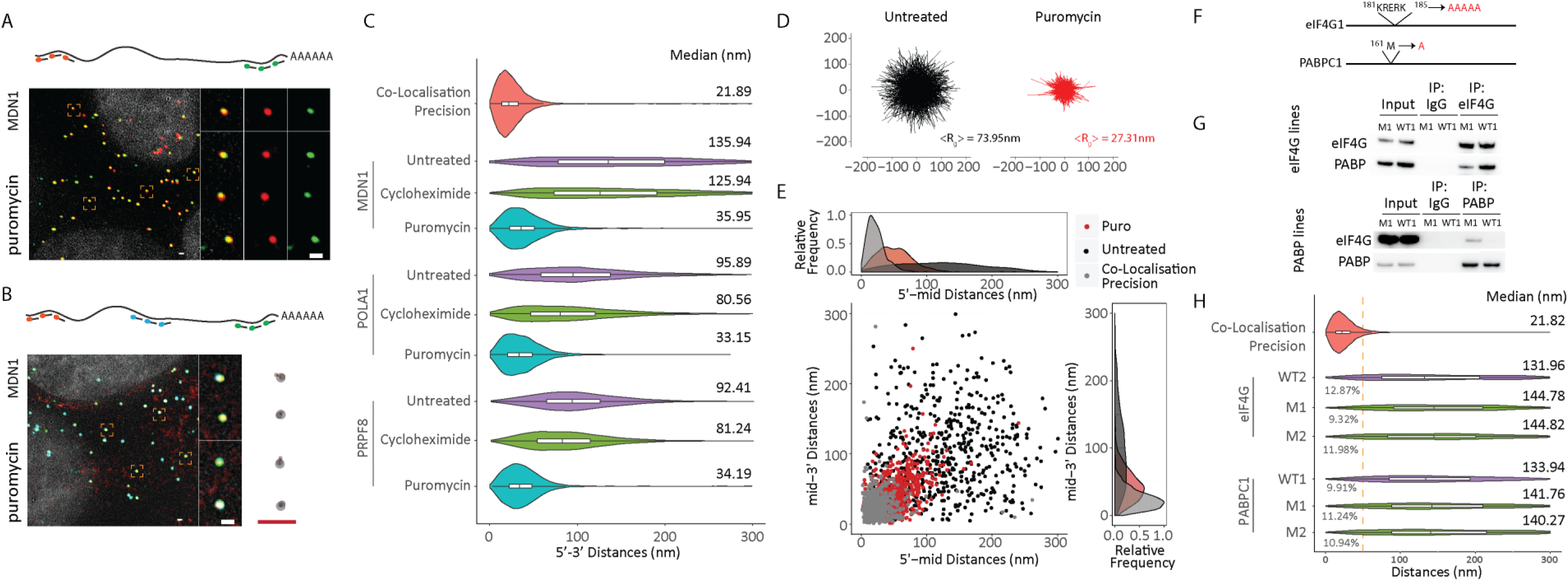
Open mRNP conformation correspond to translating mRNA. (**A, B)** 5’ and 3’ or three color MDN1 smFISH in HEK 293 cells treated with puromycin (10 min, 100 μg/ml). **(C)** Violin plots showing 5’-3’ distances for MDN1, POLA1, PRPF8 mRNAs treated with cycloheximide and puromycin. White box plot inside the violin plot shows first quartile, median and third quartile. Median distances are shown on the right. **(D)** Projections of superimposed conformations from three color MDN1 smFISH in untreated and puromycin treated cells with their centers of mass in registry, n=563. Mean Radius of gyration (<R_g_>). **(E)** Scatter plot showing 5’mid and mid-3’ distances for individual RNAs. Frequency distribution are shown on top and on the right. Scale bars, 500 nm. **(F)** Sites of amino acid substitutions in eIF4G1 and PABPC1 cell lines. **(G)** Immuno-precipitation of eIF4G1 and PABPC1 from wild-type and mutant cell lines using anti-eIF4G1 and PABPC1 anti-bodies. **(H)** Violin plots showing distance distribution of colocalization precision from 5’-3’ distances for MDN1 in wild type and mutant cell lines. White box plot inside the violin plot shows first quartile, median and third quartile. Median distances are shown on the right. WT1/WT2/M1/M2 represent different clonal cell lines.

Yet, a small fraction of MDN1 mRNAs in untreated cells showed 5’ and 3’ ends less than 50nm apart, distances compatible with mRNAs organized a closed-loop configuration (Fig. S2A). To determine whether this fraction indeed represents closed-loop conformations mediated by PABC1-eIF4G1, we constructed cell lines mutating key residues required for PABC1-eIF4G1 integrations using CRISPR/Cas9 gene editing (Fig. 2F). These cells showed reduced PABC1-eIF4G1 integrations but had minimal effects on cell survival and overall translation, with a slight increase of 80S in the eIF4G mutant cell line (Fig. 2G, S4A-F). The fraction of MDN1 mRNAs with 5’ – 3’ distances below 50 nm remained unchanged, indicating that the small 5’-3’ colocalizing fraction does not represent PABC1-eIF4G1 mediated interactions, but either non-translating mRNAs, or mRNPs where the ends are close to each other independent of a PABC1-eIF4G1 interaction, possibly due to the flexibility of the RNA polymer (Fig. 2H).

Our data demonstrates that translation causes a decompaction of mRNPs and a separation of 5’ and 3’ regions of mRNAs. If translation is required for an open RNP conformation, non-translating RNAs, such as cytoplasmic long non-coding RNAs (lncRNAs) should show a similar level of compaction than non-translating mRNAs, and, moreover, their compaction should be unaffected by translation inhibitors. We therefore measured end-to-end distances for two lncRNAs, TUG1 (6,091 nt) and OIP5-AS1 (8,300 nt), which were previously found to be present in both the nucleus and the cytoplasm (*18*). Both lncRNAs contain short putative ORFs that could lead to their association with ribosomes, however, their translation will be limited to the very 5’ of their RNA (*19*). As shown in Fig. 3A, 5’ and 3’ labeled TUG1 and OIP5-AS1 lncRNAs displayed a more compact conformation compared to the similarly sized PRPF8 mRNA. In addition, 5’-3’ distances of TUG1 and OIP5-AS1 lncRNAs were unaffected by puromycin, further suggesting that decompaction of cytoplasmic mRNAs requires the formation of polysomes (Fig. 3B). Consistent with the requirement of ribosome occupancy for RNA decompaction, 5’-to-mid region of MDN1 mRNA compacts before the mid-to-3’ region when cells are treated for 10 min with the translation inhibitor homoharringtonine (Fig. 3C,D, S5).

**Figure 3:**
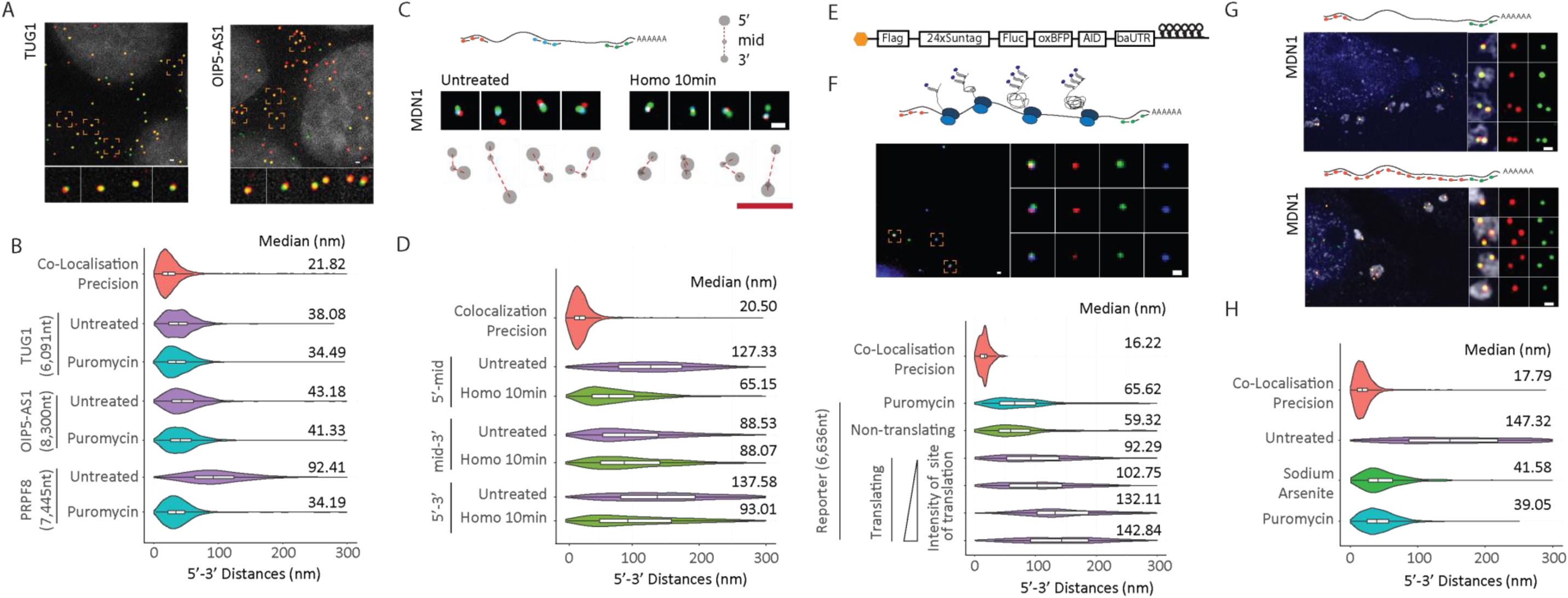
Ribosome occupancy determines mRNP compaction. **(A)** smFISH visualizing 5’ and 3’ ends of TUG1 and OIP5-AS1 lncRNAs. Nuclei are visualized by DAPI staining (grey). **(B)** Violin plots showing 5’-3’ distance distribution of cytoplasmic TUG1 and OIP5-AS1 RNAs in untreated and puromycin treated cells compared to PRPF8 mRNAs. **(C)** smFISH using 5’ (red), 3’ (green), and middle probes (cyan) respectively for untreated or homoharringtonine (100μg/ml, 10 min) treated cells and cartoon depicting different mRNA conformations. At a translation speed of 5 amino acids per second, all translating ribosomes will have reached at least the mid region of MDN1. (**D)** Violin plots showing 5’-mid, mid-3’ and 5’-3’distance distribution of cytoplasmic MDN1 mRNAs in untreated and homoharringtonine treated cells. **(E)** Cartoon depicting the SINAPs construct **(F)** Images showing 5’ and 3’ smFISH and anti-GFP immunofluorescence (top), and violin plots depicting 5’-3’ distances for puromycin treated, non-translating and translating mRNAs. Translating mRNAs were clustered in 4 groups (k-means) according to intensity of anti-GFP signal (bottom). **(G)** 5’ – 3’ or 3’ and tiling MDN1 smFISH in U2OS cells treated with arsenite (1 hour, 2 mM). Stress granules are visualized using an oligo dT probe (white). Nuclei are visualized by DAPI staining (blue). **(H)** Violin plots comparing MDN1 mRNA 5’-3’ distance distribution for untreated, arsenite and puromycin treated cells. White box plot inside the violin plot shows first quartile, median and third quartile. Median distances are shown on the right. Scale bars, 500 nm.

To directly couple translational status and 5’-to-3’ distance, we employed a reporter system developed for Single Molecule Imaging of Nascent Peptides (SINAPs), where nascent proteins are rapidly bound at the ribosome exit channel by a fluorescently labeled single-chain antibody (scFv-sfGFP) and fluorescence intensity, therefore, is proportional to the number of ribosomes on a specific mRNA (Fig. 3E) (*20, 21*). The SINAPs reporter was transfected into U2OS cells stably expressing the scFv-sfGFP fusion, cells were fixed after 24 hours and ribosome occupancy and mRNA conformation simultaneously measured by smFISH and immunofluorescence targeting scFv-sfGFP using an anti-GFP antibody. As shown in Fig. 3F, translating mRNAs show more open conformations relative to non-translating mRNAs. Moreover, the relative intensity of the translation site increases with 5’ −3’ distance, suggesting ribosome occupancy decompacts the mRNA and separates the ends. Consistently, puromycin treatment prior to fixation, eliminates translation site signals and compacts the mRNAs.

If eviction of ribosomes from translating mRNAs by puromycin results in a strong compaction of mRNA, mRNAs that are translationally repressed in response to external stimuli or environmental triggers should also acquire a compact conformation. Treatment with sodium arsenite inhibits translation through phosphorylation of eIF2α and results in disassembly of polysomes and sequestration of mRNAs in stress granules (*22, 23*). We found that upon induction of stress granule assembly in U2OS cells, following treatment with arsenite for 1 hour, cytoplasmic MDN1 mRNAs not only relocate to stress granules, but also show a highly compact conformation, as observed using 5’-3’ and tiling probes (Fig. 3G). End-to-end measurements for MDN1 mRNAs in stress-granules showed a level of compaction similar to that seen in puromycin-treated cells (Fig. 3H), and similar compaction was also observed for POLA1 and PRPF8 mRNAs under the same conditions (Fig. S6A). Interestingly, not all mRNAs accumulated in stress granules; however, most mRNAs that remained outside showed the same level of compaction as those within, suggesting that translation inhibition occurs independently of mRNA sequestration to stress granules, as previously suggested (*22*). Moreover, a fraction of TUG1 and OIP5-AS1 was also found localized to stress granules, and this localization did not alter their compaction (Fig. S6B).

We then asked whether the compacted state of mRNAs found within stress granules, or after puromycin treatment, reflects a default state for non-translating cellular mRNPs. In the nucleus, nascent mRNAs are co-transcriptionally spliced and assembled into mRNPs resulting in the binding of a large set of RBPs, including the exon-junction complex and SR proteins (*2, 24, 25*). During translation in the cytoplasm, many RBPs bound to the open reading frame are evicted by the ribosome. mRNAs that have been translated and then go into a translationally silent state might therefore be bound by fewer proteins than cytoplasmic mRNAs prior to their first round of translation or nuclear mRNPs before their export to the cytoplasm. To therefore determine whether a default compaction state exists for non-translating mRNPs, we investigated the organization of nuclear MDN1 mRNAs. Compared to MDN1 mRNAs in stress granules, many nuclear MDN1 mRNAs were found in an extended conformation, yet more compacted than translating cytoplasmic mRNAs (Fig. 4A-E). Importantly, open mRNP conformations of nuclear MDN1 were not sensitive to puromycin (10 min) or homoharringtonine (1 h) treatment (Fig. S7), suggesting that assembly of nuclear mRNPs results in more extended mRNP compared to translationally inhibited mRNPs.

**Figure 4:**
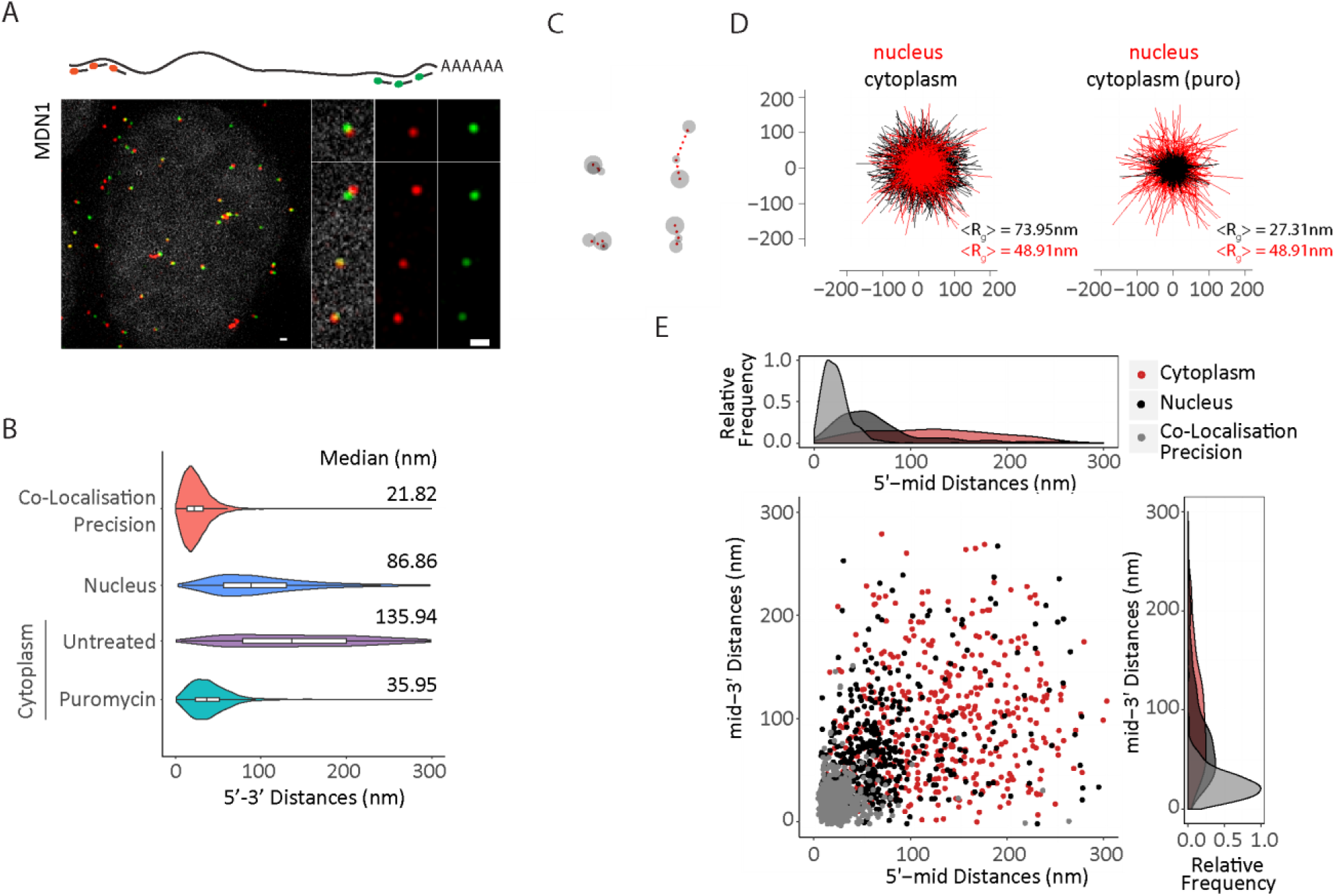
Organization of nuclear MDN1 mRNAs. **(A)** 5’-3’ MDN1 smFISH of nuclear mRNAs. The nucleus was stained with DapI (gray). **(B)** Violin plots comparing MDN1 mRNA 5’-3’ distance distribution of nuclear and cytoplasmic mRNAs. White box plot inside the violin plot shows first quartile, median and third quartile. Median distances are shown on the right. **(C)** Representative conformations of nuclear MDN1 mRNAs measured by 5, middle and 3’ labeling as in 1E. **(D)** Projections of superimposed conformations from C with their centers of mass in registry, compared to untreated or puromycin treated cytoplasmic MDN1 mRNAs, n=452. Mean Radius of gyration (<R_g_>). **(E)** Scatter plot comparing 5’-mid and mid-3’ distances for individual nuclear and cytoplasmic MDN1 mRNAs. Frequency distribution are shown on top and on the right. Scale bars, 500 nm.

Electron microscopy (EM) studies visualizing the 35 kb-long nuclear Balbiani ring mRNPs in the dipteran *Chironomus tentans* have suggested that nuclear mRNPs exist as compact particles with 5’ and 3’ ends in close proximity (*3*). Our observations do not support such a globular structure, but instead are consistent with data from recent RNA ImmunoPrecipitation and Proximity Ligation in Tandem experiments (RIPPLiT) and EM images of purified nuclear mRNPs from yeast, which suggest that mRNAs assemble in a linearly compacted polymer that can exist in various states of compaction (*4, 26*).

The current model suggests translation occurring in a closed-looped configuration is based on biochemical characterization of PABC1 and eIF4F interactions, and supported by EM images of *in vitro* assembled close-looped mRNPs (*11, 12, 16, 27, 28*). On the other hand, studies investigating polysome conformation by EM *in vivo*, have observed polysomes in various conformations with only some compatible with a closed-loop translation model (*7, 29*). However, as mRNA was not visualized in these polysomes, it was not possible to assess 5’ −3’ end mRNA interactions based on those EM images. Overall, our data demonstrates that in human cells, at least for some mRNAs, translation does not occur in a closed loop conformation. Interestingly, recent studies in the yeast *S. cerevisiae* suggest that not all mRNAs are bound to the same extent by the closed-loop factors, however, it is unclear how this relates to closed-loop RNA conformations for these mRNAs in cells, and whether this is true in higher eukaryotes (*30–32*).

Regulatory elements in mRNAs are often located within the 3’UTR and modulate processes at the 5’ end, such as de-capping or translation initiation (*13, 28*). Signal transmission from the 3’ to the 5’ likely requires a flexible RNA polymer that allows both ends to meet. However, we have little understanding of the biophysical properties of mRNPs *in vivo*. Understanding whether the open conformations observed here for mRNAs reflect continuously changing, dynamic conformations that result in frequent transient 5’-3’end interactions, and to determine if the biophysical properties of mRNAs and mRNPs allow frequent intra-molecular interactions in general, will require new tools that allow us to study mRNP organization within cells in real time.

## Acknowledgement

We thank members of the Zenklusen and Rissland laboratories, Marlene Oeffinger, Nicole Francis, Steve Michnick, Jeff Kieft and Julie Claycomb for critical discussion and comments on the manuscript. This work has been supported CIHR (Project Grant-366682), FRQ-S (Chercheur-boursier Junior 2) and CFI (DZ).

